# Leucettinib-21 decreases dosage effects of DYRK1A in human trisomy 21 iPSC-derived neural cells

**DOI:** 10.64898/2026.02.05.704014

**Authors:** Nicole R. West, Mattias F. Lindberg, Julien Dairou, Shawn MacGregor, Sahith Puthireddy, Laurent Meijer, Anita Bhattacharyya

**Affiliations:** Waisman Center, University of Wisconsin-Madison, Madison WI, USA; Cellular and Molecular Biology Graduate Program, University of Wisconsin-Madison, Madison WI, USA; Perha Pharmaceuticals, Perharidy Peninsula, 29680 Roscoff, France; Laboratoire de Chimie et Biochimie Pharmacologiques et Toxicologiques, UMR 8601 CNRS, Université Paris Cité, France; Department of Cell and Regenerative Biology, School of Medicine and Public Health, University of Wisconsin-Madison, Madison WI, USA

**Keywords:** DYRK1A, DYRK1A inhibitors, Down syndrome, Induced pluripotent stem cells (iPSCs), Cell proliferation, Tau Phosphorylation

## Abstract

Dysregulated expression and activity of DYRK1A, dual specificity tyrosine phosphorylation regulated kinase 1A, is a feature of several neurodevelopmental and neurodegenerative diseases, including Down syndrome, DYRK1A syndrome, autism spectrum disorders, Alzheimer’s disease, and Parkinson’s disease. Thus, manipulating DYRK1A activity in the brain has emerged as a potential therapeutic target for neurological disorders. Several DYRK1A inhibitors have shown promise for improving cognition in rodent models of Down syndrome and Alzheimer’s disease, for example, but the ability to affect DYRK1A levels or activity in relevant human cells has not been established. We filled this gap by testing the effects of a new DYRK1A inhibitor on trisomy 21 induced pluripotent stem cell derived neural progenitor cells and neurons, where DYRK1A expression and activity are increased. Our results demonstrate that Leucettinib-21, a potent and selective low-molecular weight pharmacological inhibitor of DYRK1A, decreases DYRK1A activity in human trisomy 21 neural progenitor cells and cortical neurons. We show for the first time that Leucettinib-21 reduces DYRK1A activity in a relevant human disease model, supporting future human trials.

**Summary Statement:** We show for the first time that Leucettinib-21, a pharmacological inhibitor of DYRK1A, decreases DYRK1A activity in a human iPSC-derived neural cell culture model of Down syndrome.

## Introduction

DYRK1A is evolutionarily conserved and belongs to the CMGC family of kinases that play a role in a wide range of cellular functions. DYRK1A autophosphorylates a tyrosine residue within its activation loop and phosphorylates serine and threonine residues on its substrates (Duchon and Herault, 2016, Atas-Ozcan et al., 2021, Rammohan et al., 2022). DYRK1A is ubiquitously expressed in organs across development and adulthood and plays a key role in maintaining normal cell proliferation, differentiation, and function (Atas-Ozcan et al., 2021, Guimera et al., 1999, Deboever et al., 2022).

### DYRK1A is dosage sensitive

Because of the pivotal role of DYRK1A in regulating cellular process, the proper dosage of DYRK1A is necessary for normal brain development and function. DYRK1A syndrome, a rare *DYRK1A* haploinsufficiency, is among the most common monogenic forms of intellectual disability (Courraud et al., 2021). DYRK1A syndrome results from microdeletions in chromosome 21q22.12q22.3, single-nucleotide variants (SNVs), translocations, or small insertions or deletions in the gene (Ji et al., 2015). Microcephaly, intellectual disability, autism spectrum disorder (ASD), speech impairment, seizures, skeletal and gait abnormalities, and eye defects are characteristic features of DYRK1A syndrome (Ji et al., 2015).

DYRK1A overexpression has been linked to several cancers, including acute leukemia, glioblastoma, head and neck squamous cell carcinoma, hepatocellular carcinoma, and pancreatic ductal adenocarcinoma due to its role in cell cycle progression, DNA damage repair, and cancer stem cell maintenance (Rammohan et al., 2022, Fernández-Martínez et al., 2015, Boni et al., 2020). DYRK1A promotes tumor growth in some cellular contexts and DYRK1A overexpression correlates with poor prognosis in certain cancers (Laham et al., 2022, Laham et al., 2024, Li et al., 2019).

*DYRK1A* is encoded on chromosome 21 (Hsa21) and is overexpressed in Down syndrome (DS, trisomy 21, T21) that is caused by the triplication of Hsa21 (Lejeune et al., 1959b, Lejeune et al., 1959a). DS is the leading genetic cause of intellectual disability with IQs ranging from mild to severe and an average IQ around 50 (Mégarbané et al., 2013, Kłosowska et al., 2022, Lejeune et al., 1959a, Lejeune et al., 1959b, Hassold and Hunt, 2001, Fidler and Nadel, 2007, Hattori et al., 2000). In addition to the cognitive deficits, individuals with DS have increased risk for congenital heart defects, leukemia, gastrointestinal problems, vision and hearing impairment, and early onset of Alzheimer’s disease (AD) (Roizen, 2010, Capone et al., 2018, Lott and Head, 2019). The role of DYRK1A overexpression in the cellular processes that regulate these diverse organ defects has been an area of intense research and potential therapeutics.

### Manipulating DYRK1A expression and activity

Like many kinases, DYRK1A’s role in proliferation has made it a therapeutic target for cancer and other diseases, including diabetes (Rammohan et al., 2022, Barzowska et al., 2021, Deboever et al., 2022). Several DYRK1A inhibitors exist including Epigallocatechin gallate (EGCG), Harmine, Leucettine L41, Leucettinib-21 (LCTB-21), Lorecivivint, Cirtuvivint, PST-001, and others (Lindberg et al., 2023a, Lindberg et al., 2023b). LCTB-21, a potent, low-molecular weight DYRK1A inhibitor developed by Perha Pharmaceuticals, improves cognitive function in rodent models of DS and AD (Lindberg et al., 2023b, Deau et al., 2023), and it has passed preclinical regulatory safety trials in rats and minipigs (Meijer et al., 2024).

LCTB-21 is currently in a Phase 1 clinical trial to investigate its pharmacokinetics, safety and tolerability in healthy volunteers, followed by evaluation in a small cohort of individuals with DS and patients with AD (NCT06206824) (Meijer et al., 2024). However, this drug candidate has not yet been evaluated in a human DS model, which could provide insight into LCTB-21’s mechanism of action. Utilizing trisomy 21 induced pluripotent stem cells (iPSCs) and isogenic controls (Giffin-Rao et al., 2022), we tested the efficacy of LCTB-21 in human iPSC-derived neural cells to validate its effectiveness in a human disease model. The kinase-inactive isomer iso-Leucettinib-21 (iso-LCTB-21) was used as a negative control. Our results demonstrate that LCTB-21 reduces DYRK1A activity and subsequent phosphorylation of DYRK1A targets in human neural cells. This work establishes a disease-relevant cellular model that can be used in future studies to delineate LCTB-21’s mechanism of action.

## Results

### Leucettinib-21 inhibits the activity of DYRK1A in human neural progenitor cells

Because DYRK1A plays a role in regulating cell cycle progression, we tested the efficacy of LCTB-21 in proliferating neural progenitor cells. T21 and isogenic control (euploid) iPSCs were differentiated into neural progenitor cells (NPCs) and treated with DMSO, LCTB-21, or iso-LCTB-21 (Figure 1A) once neural rosettes formed at Day 18 (Hunt et al., 2019, Fedorova et al., 2019) (Figure 1B). Control and T21 cultures had similar enriched proportions of PAX6+ and SOX2+ NPCs and treatment with LCTB-21 or iso-LCTB-21 did not affect the cellular composition compared to vehicle (DMSO) (Supporting Figures 1A-1C). To determine if the drug candidate affects NPC viability, we used the CellTiter-Glo 2.0 Assay (Promega) to assess viability and found that 0.01 and 0.1 μM LCTB-21 treatment has no effect on the viability of NPCs relative to DMSO. A higher dose (1 μM) of LCTB-21 is toxic to ∼20% of control and ∼25% of T21 NPCs (Supporting Figure 1D). As previously reported (Martí et al., 2003), DYRK1A protein is expressed in the cytoplasm and the nucleus and is localized similarly in control and T21 NPCs (Supporting Figure 1E).

**Figure 1.**
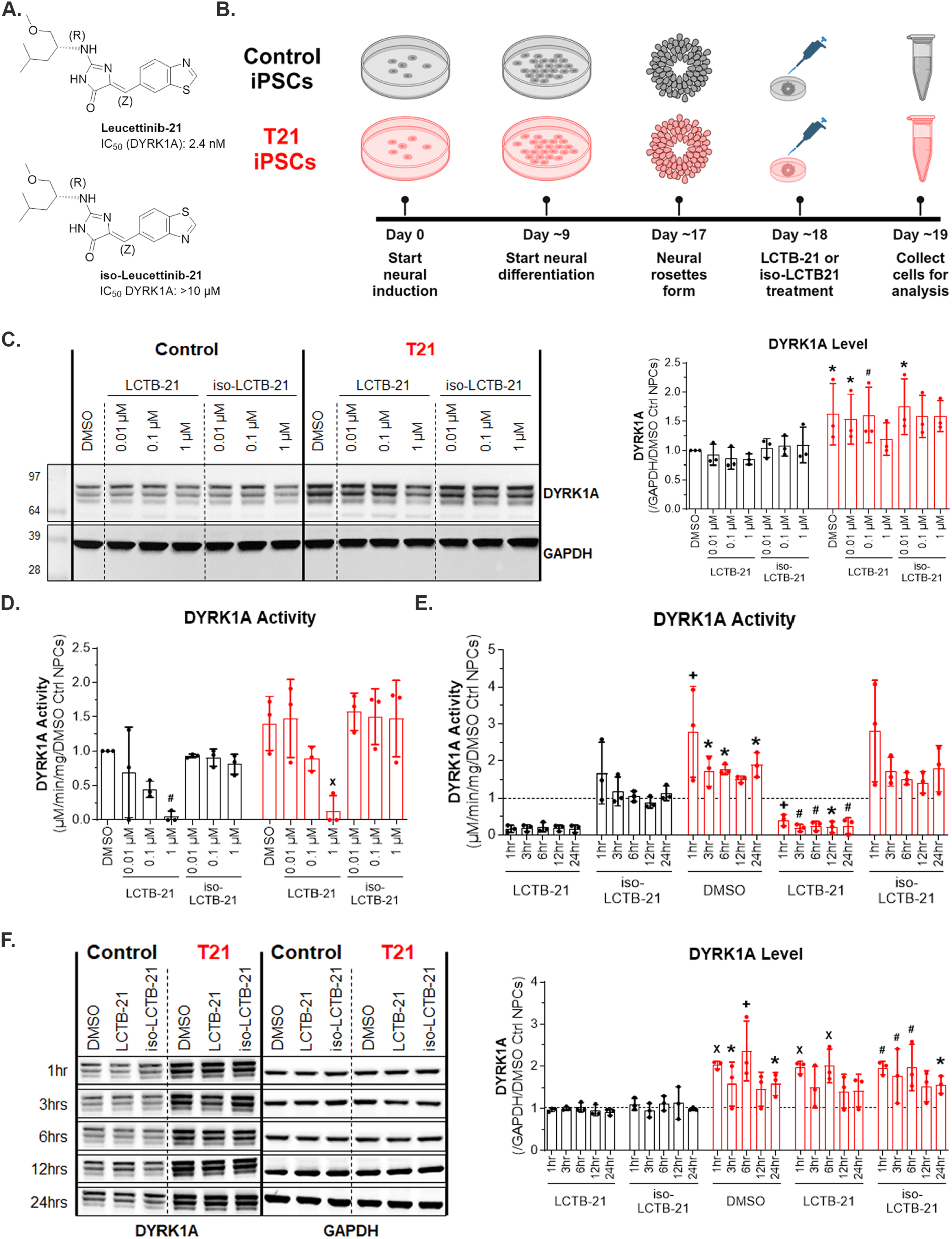
LCTB-21 reduces the activity of DYRK1A in human control and T21 neural progenitor cells. A) Structure of Leucettinib-21 and its kinase-inactive isomer, iso-Leucettinib-21. IC_50_ values on DYRK1A are provided. B) Schematic of iPSC-derived NPC differentiation and treatment (Created with BioRender.com). C) Western blot and quantification of DYRK1A levels in control and T21 NPCs. DYRK1A protein is increased in T21 NPCs, and treatment with LCTB-21 does not affect the protein levels of DYRK1A. D) Quantification of DYRK1A activity in NPCs. DYRK1A activity is increased in T21 NPCs. DYRK1A activity is decreased in a dose-dependent manner in control and T21 NPCs when treated with increasing concentrations of LCTB-21. E) Quantification of DYRK1A activity in NPCs over 24 hours. DYRK1A activity of DMSO treated T21 NPCs is increased relative to DMSO treated control NPCs (represented by the dashed line) at the respective time. DYRK1A activity in T21 NPCs is decreased within 1 hour and persists for at least 24 hours when treated with 1 μM LCTB-21 compared to DMSO treated T21 NPCs at the respective time. F) Western blot and quantification of DYRK1A levels in control and T21 NPCs over time. DYRK1A is increased in T21 NPCs compared to control NPCs at the respective treatment and time. DMSO treated control NPCs are represented by the dashed line. Treatment with LCTB-21 does not affect the levels of DYRK1A. Data presented as mean + SD. P-values from Two-way ANOVA with post-hoc tests are presented as * ≤ 0.05, # ≤ 0.01, X ≤ 0.001, + ≤ 0.0001. Two-way ANOVA results, post-hoc tests, and P-values, and excluded values are listed in Supporting Table 3.

We assessed the expression of DYRK1A protein in iPSC-derived NPCs. As expected, DYRK1A protein levels in T21 NPCs are increased relative to control (Figure 1C). Protein levels of DYRK1A do not change significantly in control or T21 NPCs following the addition of LCTB-21 or iso-LCTB-21 compared to the DMSO control. However, there is a slight decrease in the DYRK1A level in T21 NPCs treated with 1 μM LCTB-21 (Figure 1C). LCTB-21 inhibition of DYRK1A activity at the 1 μM dose may prevent DYRK1A autophosphorylation at Ser97, leading to its degradation (Kii et al., 2016).

Consistent with DYRK1A protein expression, DYRK1A kinase activity is increased ∼1.5-fold in T21 NPCs compared to control. Treatment with LCTB-21 decreases DYRK1A activity in a dose-dependent manner in both control and T21 NPCs, consistent with observations in HT-22 hippocampal mouse cells (Lindberg et al., 2023b) (Figure 1D). To determine the time course of LCTB-21 action on DYRK1A kinase activity, DYRK1A kinase activity was measured in NPCs treated with 1 μM LCTB-21 at 1 hour, 3 hours, 6 hours, 12 hours, and 24 hours (Figure 1E). DYRK1A activity is reduced by more than 80% in control and T21 NPCs within 1 hour of LCTB-21 treatment, and reduction of DYRK1A activity persists for at least 24 hours after LCTB-21 treatment. Consistent with the dose-response assay (Figure 1C), increased DYRK1A protein level is detected in T21 NPCs. LCTB-21 and iso-LCTB-21 treatments do not affect DYRK1A protein levels (Figure 1F). These results demonstrate that LCTB-21 is effective at inhibiting the activity of DYRK1A in human NPCs.

### DYRK1A inhibition decreases phosphorylation of the DYRK1A target, cyclin D1, in human neural progenitor cells

Cyclin D1 is phosphorylated by DYRK1A at Thr286 (pT286-cyclin D1) (Soppa et al., 2014, Ashford et al., 2013) and plays an integral role in the expression of cell cycle genes, including *MKI67* (Hille et al., 2016), that can affect cell cycle progression (Rammohan et al., 2022, Chen et al., 2013, Laham et al., 2024). DYRK1A imbalance leads to altered cell cycle progression in mouse and cell models of DS (Chakrabarti et al., 2007, Sharma et al., 2022, Najas et al., 2015) (Figure 2A). Since LCTB-21 reduces the activity of DYRK1A in NPCs, we asked whether LCTB-21 has downstream effects on the phosphorylation of the DYRK1A target, cyclin D1.

**Figure 2.**
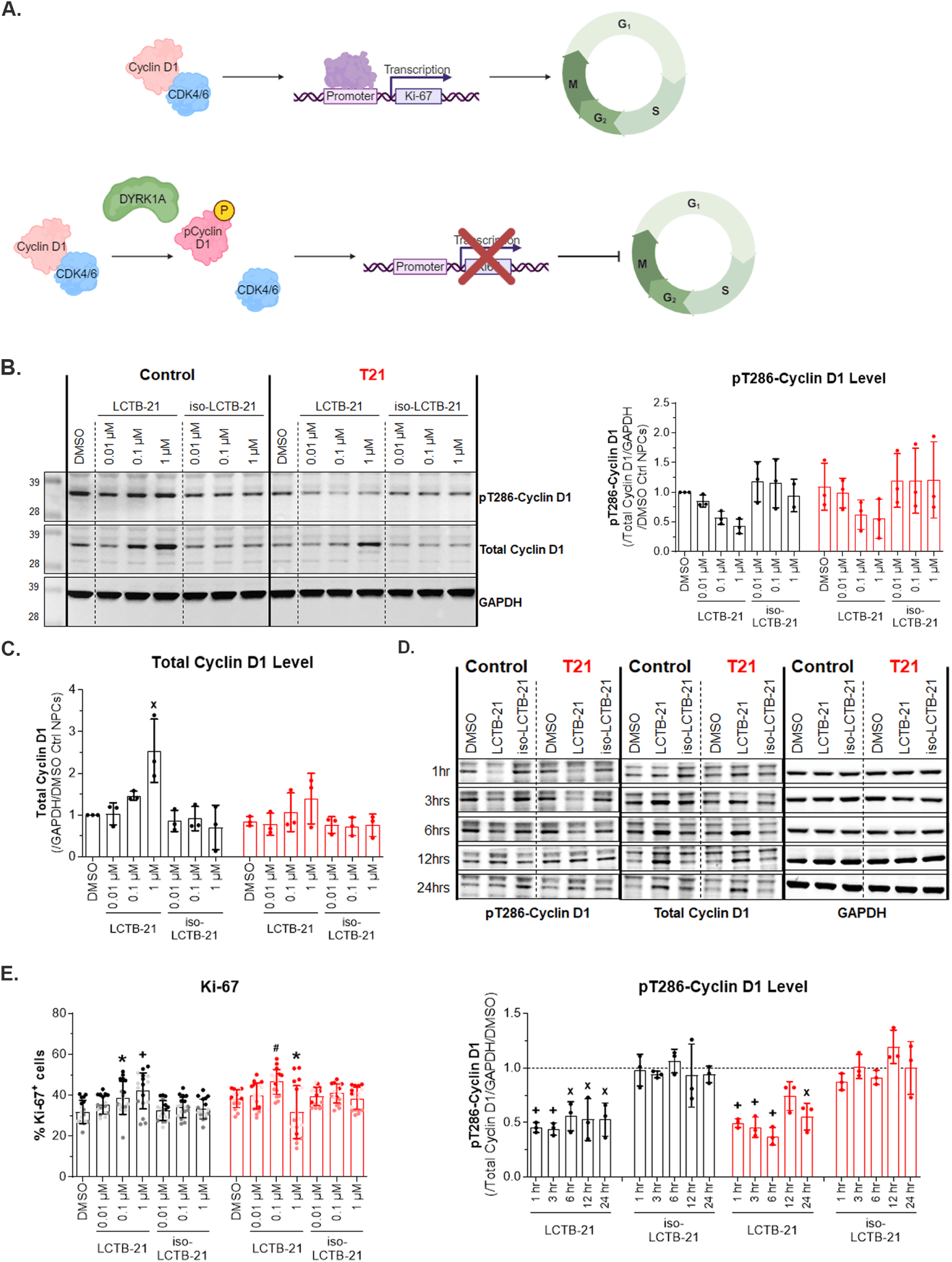
DYRK1A inhibition decreases phosphorylation of the DYRK1A target, cyclin D1, in human neural progenitor cells. A) Schematic of DYRK1A and cyclin D1 phosphorylation (Created with BioRender.com). When cyclin D1 is phosphorylated by DYRK1A, downstream cell cycle genes, including *MKI67*, are not transcribed. B) Western blot and quantification of cyclin D1 and pT286-cyclin D1 levels in control and T21 NPCs. pT286-cyclin D1 levels decrease as DYRK1A activity decreases with increasing doses of LCTB-21. C) Cyclin D1 levels increase with increasing doses of LCTB-21. D) Western blot and quantification of cyclin D1 and pT286-cyclin D1 levels in control and T21 NPCs over time. pT286-cyclin D1 is decreased in control and T21 NPCs within 1 hour and persists for at least 24 hours when treated with 1 μM LCTB-21 relative to respective DMSO (represented by the dashed line). E) Ki-67+ cells in the cell cycle. More T21 NPCs are Ki-67+, and with the exception of T21 NPCs treated with 1 μM LCTB-21, the number of Ki-67+ cells increase with LCTB-21 treatment. Data presented as mean + SD. P-values from Two-way ANOVA with post-hoc tests are presented as * ≤ 0.05, # ≤ 0.01, X ≤ 0.001, + ≤ 0.0001. Two-way ANOVA results, post-hoc tests, and P-values, and excluded values are listed in Supporting Table 3.

We found a dose-dependent reduction in the phosphorylation of cyclin D1 at Thr286 relative to total cyclin D1 in both control and T21 NPCs when treated with LCTB-21 (Figure 2B), concomitant with the dose-dependent decrease of DYRK1A activity (Figure 1D). Thr286-phosphorylated cyclin D1 is unstable, and dephosphorylated cyclin D1 shows increased stability (Diehl et al., 1997). DYRK1A inhibition by LCTB-21 thus leads to cyclin D1 accumulation in both control and T21 NPCs (Figure 2C).

To determine how quickly LCTB-21 and the reduction in DYRK1A activity are affecting cyclin D1 phosphorylation, we assessed pT286-cyclin D1 levels relative to total cyclin D1 over time in NPCs treated with 1 μM LCTB-21 (Figure 2D). pT286-cyclin D1 is reduced in control and T21 NPCs within 1 hour of treatment with LCTB-21 and this reduction persists for 6 hours (Supporting Figure 1F). Concomitant with the decrease in T286 phosphorylation, cyclin D1 accumulates between 6 and 24 hours post LCTB-21 treatment (Supporting Figure 1G). Overall, inhibition of DYRK1A by LCTB-21 results in a reduction in the ratio of pT286-cyclinD1 to total cyclin D1, which is maintained in a similar manner from 1 hour to 24 hours (Figure 2D).

We next asked whether reduction of DYRK1A activity alters the proliferation of NPCs. We analyzed the number of cycling (Ki-67+) cells upon LCTB-21 treatment (Figure 2E). The number of Ki-67+ cells slightly increases in proportion to LCTB-21 dosage in control NPCs. More T21 NPCs are cycling upon LCTB-21 treatment. The trend is the same in T21 NPCs with the exception of the 1 µM dose at which T21 Ki-67+ cells decrease is likely due to toxicity (Supporting Figure 1A). DYRK1A inhibition leads to a reduction in pT286-cyclin D1 and an accumulation of cyclin D1, favoring an increase in Ki-67+ NPCs in the cell cycle.

### Leucettinib-21 inhibits DYRK1A activity in human cortical neurons in a dose-dependent manner

Improper *DYRK1A* dosage is associated with intellectual disability, thus it is important to understand how DYRK1A inhibition affects neuronal function. We tested the efficacy of LCTB-21 in iPSC-derived cortical neurons. Neural progenitor cells were differentiated into TBR1+ cortical neurons (Supporting Figure 2A) and treated with DMSO, LCTB-21, or iso-LCTB-21 (Figure 3A). 24-hour treatment with LCTB-21 has no effect on viability of 4-week cortical neurons relative to DMSO treated controls, but there was a slight decrease in viability of T21 cortical neurons treated with iso-LCTB-21 (Figure 3B).

**Figure 3.**
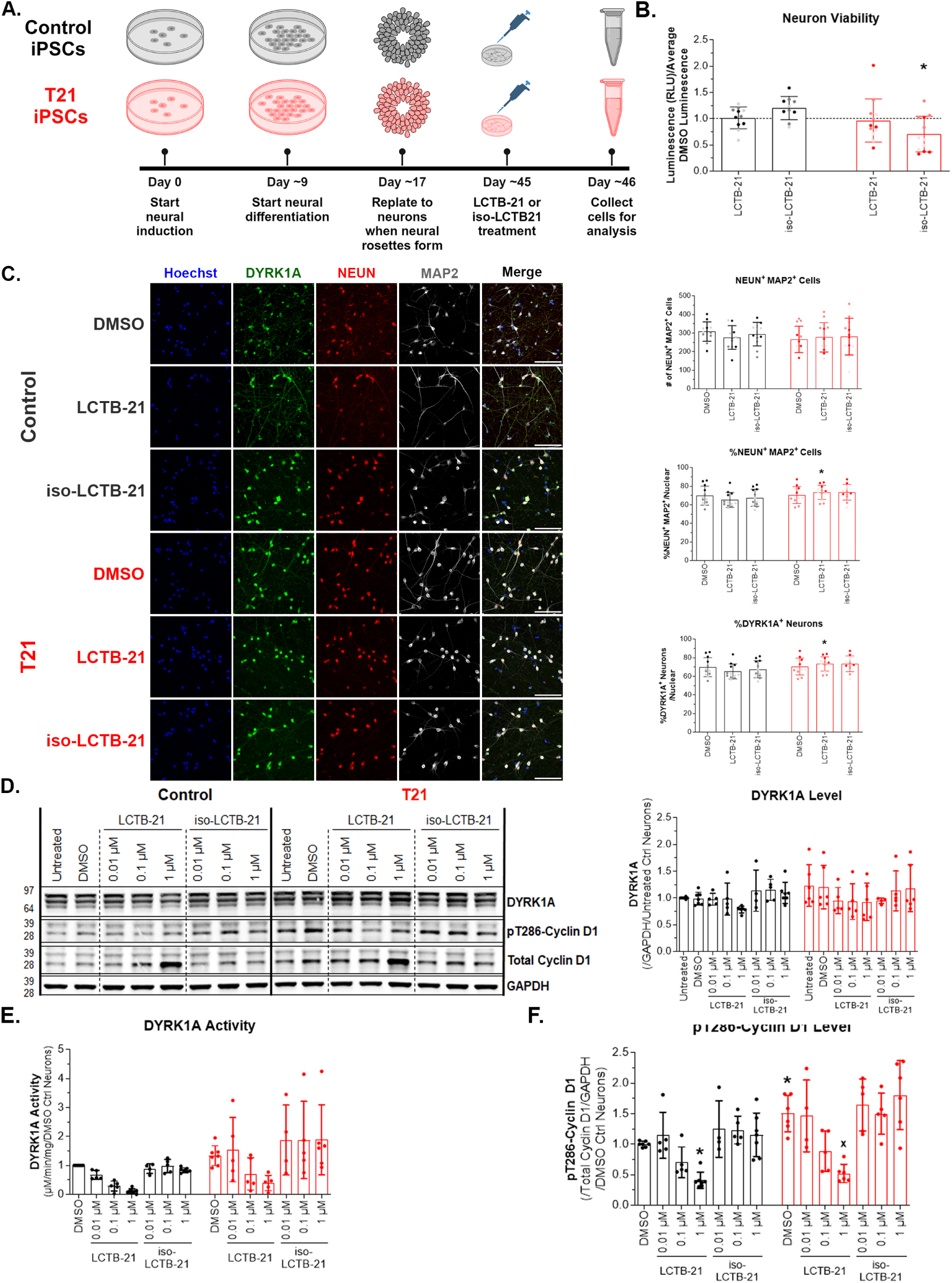
LCTB-21 inhibits DYRK1A activity in human cortical neurons in a dose-dependent manner. A) Schematic of iPSC-derived cortical neuron differentiation and treatment (Created with BioRender.com). B) Viability quantification of control and T21 neurons treated with LCTB-21 and iso-LCTB-21. LCTB-21 treatment has no effect on viability of 4-week cortical neurons compared to respective DMSO (represented by the dashed line). C) Representative images and quantification of cortical neuron cultures (40x objective, 100 μm scale bar). LCTB-21 treatment has no effect on the percentage of cortical neurons compared to respective DMSO. In the LCTB-21 treatment, there is a slight increase in the number of T21 cortical neurons compared to the control. D) Western blot and quantification of DYRK1A levels in control and T21 cortical neurons. DYRK1A levels are the same in T21 and control neurons. Treatment with LCTB-21 does not affect the levels of DYRK1A. E) Quantification of DYRK1A activity in cortical neurons. DYRK1A activity is increased in T21 cortical neurons. DYRK1A activity is decreased in a dose-dependent manner in control and T21 cortical neurons when treated with increasing concentrations of LCTB-21. F) Western blot quantification of pT286-cyclin D1 and cyclin D1 levels in control and T21 cortical neurons. pT286-cyclin D1 levels decrease in a dose-dependent manner with LCTB-21 treatment. pT286-cyclin D1 levels in T21 neurons are higher relative to control neurons. pT286-cyclin D1 levels decrease in both T21 and control neurons treated with 1 μM LCTB-21 relative to their respective DMSO treatment. Data presented as mean + SD. P-values from Two-way ANOVA with post-hoc tests are presented as * ≤ 0.05, # ≤ 0.01, X ≤ 0.001, + ≤ 0.0001. Two-way ANOVA results, post-hoc tests, and P-values, and excluded values are listed in Supporting Table 3.

Control and T21 iPSC-derived cultures have similar populations of neurons (Figure 3C). Treatment with LCTB-21 has no effect on the neuron composition relative to DMSO treatment (Figure 3C). The T21 cultures treated with LCTB-21 have a slightly higher percentage of neurons compared to control cultures (Figure 3C). Cultures contained ∼70% NEUN+ MAP2+ neurons, and all neurons expressed DYRK1A (Figure 3C).

DYRK1A protein levels are not significantly higher in T21 cortical neurons compared to controls (Figure 3D). Nevertheless, consistent with the NPC data, DYRK1A activity is higher in T21 neurons (Figure 3E). There is a dose-dependent decrease in DYRK1A activity following LCTB-21 treatment (Figure 3E) in both control and T21 neurons. In postmitotic neurons, cyclin D1 plays a role in neuronal signaling and survival (Pedraza et al., 2023, Sumrejkanchanakij et al., 2003). pT286-cyclin D1 is higher in T21 cortical neurons relative to control cortical neurons (Figure 3F). Consistent with the decrease in DYRK1A activity with LCTB-21 treatment, pT286-cyclin D1 is decreased in a dose-dependent manner in T21 and control neurons treated with LCTB-21 (Figure 3F). Thus, LCTB-21 reduces DYRK1A activity in a dose-dependent manner in iPSC-derived cortical neurons, whereas iso-LCTB-21 has no effect.

### DYRK1A inhibition decreases phosphorylation of the DYRK1A target, Tau, in human cortical neurons

In neurons, DYRK1A phosphorylates diverse target proteins associated with differentiation and neuronal maintenance (Atas-Ozcan et al., 2021, Wang et al., 2017) The microtubule-associated protein, Tau, that helps maintain cellular structure and supports axonal transport (Tabeshmehr and Eftekharpour, 2023), is phosphorylated by DYRK1A at several sites, Thr212 being the best characterized site – phosphorylation at this site primes for further phosphorylation by GSK3 at Ser208 (Ryoo et al., 2007, Kimura et al., 2007). Tau phosphorylation inhibits microtubule assembly and causes Tau to aggregate into neurofibrillary tangles (NFTs) (Parra Bravo et al., 2024, Aranda-Abreu et al., 2025) (Figure 4A). Since LCTB-21 inhibits the activity of DYRK1A in cortical neurons, we tested whether LCTB-21 reduces the phosphorylation of the DYRK1A target, Tau.

**Figure 4.**
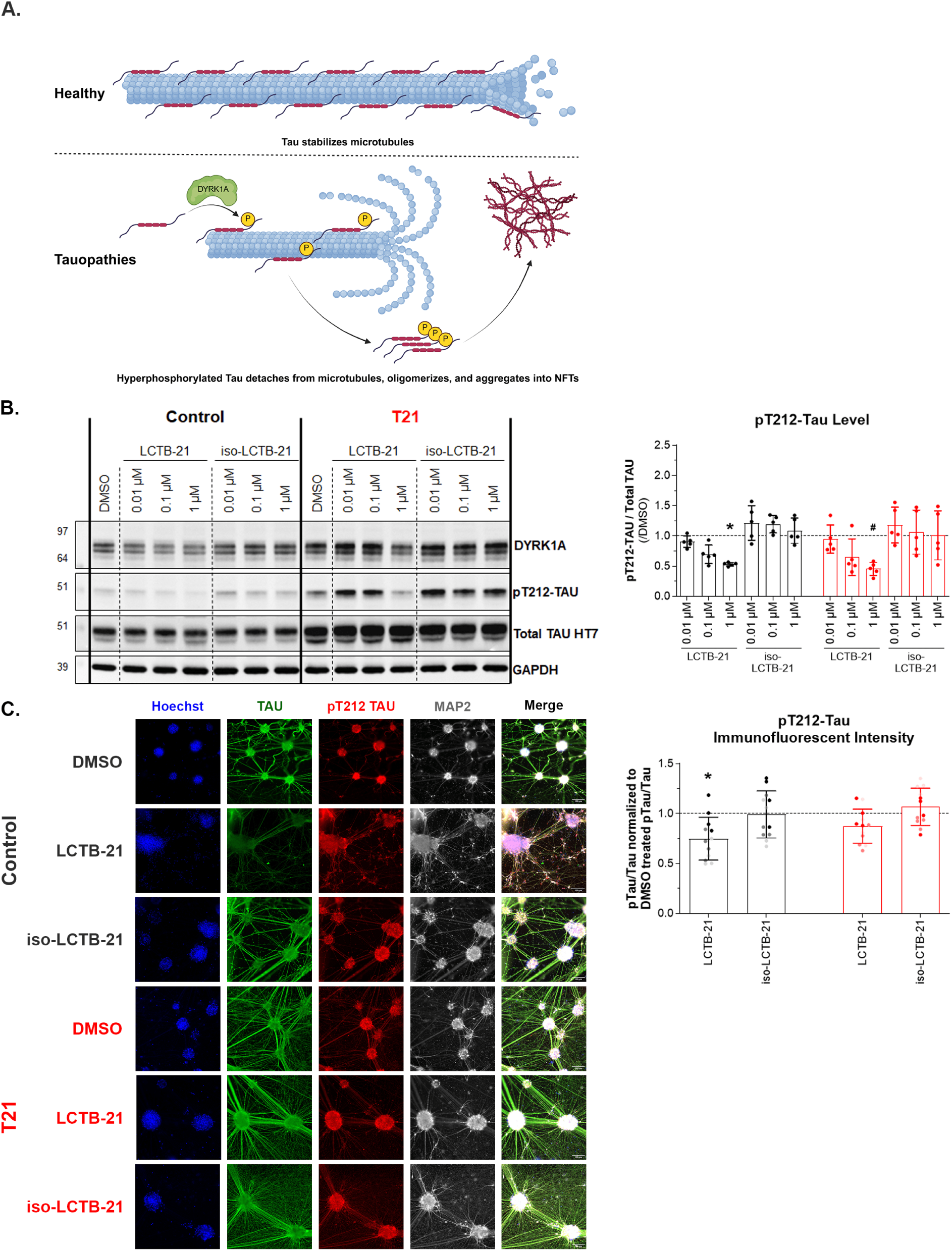
DYRK1A inhibition decreases phosphorylation of the DYRK1A target, Tau, in human cortical neurons. A) Schematic of DYRK1A and Tau phosphorylation (Created with BioRender.com). B) Western blot and quantification of Total Tau and pT212 Tau levels in control and T21 cortical neurons. pT212 Tau levels decrease as DYRK1A activity decreases with increasing doses of LCTB-21. C) Representative images (20x objective, 100 μm scale bar) and immunofluorescent intensity of pT212 Tau to Total Tau normalized to the pT212 Tau/Total Tau ratio of the respective DMSO treatment (represented by the dashed line). Data presented as mean + SD. P-values from Two-way ANOVA with post-hoc tests are presented as * ≤ 0.05, # ≤ 0.01, X ≤ 0.001, + ≤ 0.0001. Two-way ANOVA results, post-hoc tests, and P-values, and excluded values are listed in Supporting Table 3.

We found a dose-dependent decrease in the phosphorylation of Tau at Thr212 relative to total Tau in 4-week iPSC-derived cortical neurons treated with LCTB-21 for 24 hours (Figure 4B). The inactive isomer iso-LCTB-21 did not affect levels of pT212 Tau (Figure 4B). Consistent with the immunoblot results, immunofluorescence intensity for pT212 Tau relative to total Tau decreased following treatment with 1 µM LCTB-21 (Figure 4C). This reduction was significant in control neurons, while only a slight decrease was observed in the T21 neurons relative to pT212 Tau/total Tau intensity in the respective DMSO treated samples.

LCTB-21 treatment also indirectly leads to a decrease in the phosphorylation of T217 Tau (Supporting Figure 2B). DYRK1A-mediated phosphorylation of Tau primes Tau to be phosphorylated by other kinases at nearby residues (Liu et al., 2006, Cho and Johnson, 2003). As DYRK1A activity (Figure 3E) and pT212 Tau levels (Figure 4B) decrease with increasing dose of LCTB-21, pT217 levels decrease in control and T21 neurons (Supporting Figure 2B).

## Discussion

*DYRK1A* is a dosage-sensitive kinase encoded on Hsa21. It is ubiquitously expressed across development in several tissues, including the brain. DYRK1A regulates cellular processes such as cell cycle progression, neurogenesis, neuronal differentiation, axonal transport and synaptic plasticity. Dosage imbalance of *DYRK1A* and altered activity are associated with several neurodevelopmental disorders and neurodegenerative diseases. *DYRK1A* haploinsufficiency causes the neurodevelopmental disorder DYRK1A syndrome whereas overexpression contributes to DS pathology.

Due to its role in cognitive function, a potent, low-molecular weight DYRK1A inhibitor, LCTB-21, has been developed to alleviate cognitive deficits associated with DS and AD (Feki and Hibaoui, 2018, Duchon and Herault, 2016, Atas-Ozcan et al., 2021). While LCTB-21 has been shown to improve cognitive function in rodent models of DS and AD and is currently in Phase 1 clinical trial, this is the first study to test LCTB-21 in a human DS cell model. This work demonstrates that LCTB-21 effectively inhibits DYRK1A activity in an iPSC-derived neural model of DS, establishing a platform for future investigations to delineate the mechanism of action of pharmacological DYRK1A inhibitors.

LCTB-21 reduces the activity of DYRK1A without affecting DYRK1A protein levels in human iPSC-derived NPCs. At baseline, DYRK1A activity is increased in T21 NPCs. The activity of DYRK1A decreases in a dose-dependent manner in control and T21 NPCs when treated with LCTB-21. DYRK1A activity is decreased within an hour of LCTB-21 treatment and the reduction persists for at least 24 hours after treatment. The inactive isomer, iso-LCTB-21, has no effect on the activity of DYRK1A.

Inhibition of DYRK1A activity via LCTB-21 has downstream effects in human NPCs, including phosphorylation of cyclin D1 at Thr286. pT286-cyclin D1 is reduced in both control and T21 cells upon LCTB-21 treatment, concomitant with an increase in Ki-67+. This result contradicts what we expected – phosphorylation of cyclin D1 by DYRK1A suppresses cell cycle genes and ultimately proliferation. When cyclin D1 is phosphorylated by DYRK1A, the transcription factor E2F remains bound and cannot bind to promoter regions, thus suppressing expression of cell cycle genes (Yang et al., 2006), including *MKI67*. Increased expression of DYRK1A thus reduces the expression of cell cycle genes and ultimately proliferation by phosphorylating cyclin D1. The accumulation of cyclin D1 accompanied by decreased degradation of pT286-cyclin D1 raises the possibility that Ki-67+ control and T21 NPCs are accumulating in the G_1_ phase and not progressing through the cell cycle when treated with LCTB-21. Future work is needed to determine if and how LCTB-21-mediated DYRK1A inhibition affects cell proliferation.

LCTB-21 reduces the catalytic activity of DYRK1A in human iPSC-derived cortical neurons. The activity of DYRK1A decreases in a dose-dependent manner in control and T21 neurons when treated with LCTB-21 while the inactive isomer, iso-LCTB-21, has no effect on the activity of DYRK1A. The inhibition of DYRK1A activity in cortical neurons by LCTB-21 causes a decrease in phosphorylation of downstream targets cyclin D1 at Thr286 and Tau at Thr212.

The microtubule-associated protein, Tau, stabilizes microtubules providing support for axonal transportation and cytoskeletal structure. Tau phosphorylation is a dynamic process allowing for reorganization of the cytoskeleton. However, when there is an imbalance in this process and Tau becomes hyperphosphorylated, it cannot bind as readily to microtubules. Hyperphosphorylated Tau is prone to misfolding and aggregates into insoluble neurofibrillary tangles, which impairs neuron function (Boutajangout et al., 2011, Lasagna-Reeves et al., 2011).

Hyperphosphorylated Tau and neurofibrillary tangles are common pathology in several neurodegenerative disease, including AD and DS-AD (Zhang et al., 2024, Sheppard et al., 2012). In these diseases, Tau can be hyperphosphorylated at several sites, including Thr212 (Miao et al., 2019, Alonso et al., 2010) by kinases including DYRK1A (Ryoo et al., 2007). Consistent with other reports of DYRK1A inhibition reducing phosphorylation of Tau (Shukla et al., 2023, Chen et al., 2024, Tu et al., 2024, Frost et al., 2011), we show that treatment with DYRK1A inhibitor, LCTB-21, reduces the phosphorylation of Tau at Thr212 which may indirectly lead to a reduction in the phosphorylation at Thr217 by other kinases. This reduction of pT212 Tau demonstrates LCTB-21’s potential to limit Tau hyperphosphorylation and aggregation, supporting its therapeutic use for mitigating Tau pathology associated with AD, DS-AD, and other tauopathies.

Taken together, our results demonstrate that LCTB-21 decreases DYRK1A activity in human trisomy 21 neural progenitor cells and cortical neurons. Downstream targets of DYRK1A are also affected by LCTB-21 treatment. We show for the first time that LCTB-21 decreases DYRK1A dosage effects in a relevant human disease model, supporting future human trials.

## Materials and Methods

### Drug Synthesis

Leucettinib-21 (LCTB-21) and iso-Leucettinib-21 (iso-LCTB-21) were developed and synthetized by Perha Pharmaceuticals as previously described (Lindberg et al., 2023b, Deau et al., 2023).

### Statistical Analysis

Results are from at least three technical replicates from separate stem cell differentiations (batches). Data was analyzed with two-way ANOVAs and appropriate post-hoc corrections in GraphPad Prism. Data are presented as mean + SD. P-values from Two-way ANOVA with post-hoc tests are presented as * ≤ 0.05, # ≤ 0.01, X ≤ 0.001, + ≤ 0.0001. Outliers were identified with Grubbs’ Test (Alpha = 0.01) in Prism and excluded from the analysis. Two-way ANOVA results, post-hoc tests, and P-values, and excluded values are listed in Supporting Table 3.

### Cell Culture

Isogenic control (WC-24-02-DS-B) and T21 (WC-24-02-DS-M) iPSCs were established in the Bhattacharyya lab (Giffin-Rao et al., 2022) (WiCell, Madison, WI). We verified cell quality by mycoplasma testing performed routinely throughout the duration of experiments and karyotyping of each line at the conclusion of experiments to ensure karyotypes remained as expected (WiCell, Madison, WI). iPSCs were cultured at 37°C/5% CO_2_ in 6-well plates on mouse embryonic fibroblasts (MEFs) (WiCell, Madison, WI) with daily changes of hESC media (Supporting Table 1). iPSCs were passaged weekly onto fresh MEFs using 1 mg/mL Collagenase Type IV (Gibco) in DMEM/F12. At the start of neural induction on Day 0, iPSCs were treated with 10 μM SB431542 (Peprotech), 0.1 μM LDN-193189 2HCl (Selleckchem), and 2 μM XAV-939 (TOCRIS) in Neural Induction Media (NIM). At Day 8 or 9, cells were replated at a 1:1 ratio from MEFs to 6-well Matrigel-coated plates. Cells were dissociated with TrypLE Express (Gibco) at 37°C for 5 minutes. TrypLE Express was removed and 2 mL Neurobasal was added to each well. Cells were gently removed from the well by pipetting. Cells were transferred to a 15 mL tube and centrifuged at 300g for 2 minutes. Cells were resuspended in 2 mL/well of Neural Progenitor Cell (NPC) media (Supporting Table 1) with 5 μM Y-27632 dihydrochloride (TOCRIS). NPC media was changed daily.

### Treatment of iPSC derived Neural Progenitor Cells (NPCs)

Once NPCs formed neural rosettes, they were treated with DMSO, LCTB-21, or iso-LCTB-21 in NPC media (Supporting Table 1) for the indicated time. For western blot and kinase activity analysis: media was removed and 1x dPBS was added to each well. Cells were scraped off of the well and transferred to an Eppendorf tube. Cells were pelleted at 4500 rpm for 4 minutes, supernatant was removed, and cell pellets were stored at -80 °C until analysis. For immunocytochemistry: media was removed and cells were fixed in 4% PFA for 15 minutes. NPCs were washed and stored in 1x PBS until analysis.

### Treatment of iPSC derived Neurons

Once neural progenitor cells (NPCs) formed neural rosettes (Hunt et al., 2019, Fedorova et al., 2019), cells were dissociated with Accutase (Millipore Sigma) at 37^C^ for 5 minutes. Single cells were transferred to a 15 mL tube and centrifuged at 300g for 2 minutes. Supernatant was removed and cells were resuspended in NPC media. Cells were plated with 5 μM Y-27632 dihydrochloride and 200 nM γ-Secretase Inhibitor XXI, Compound E (Millipore Sigma). The following day, half of the media was replaced with Neural Differentiation Media (NDM) (Supporting Table 1) with 200 nM γ-Secretase Inhibitor XXI, Compound E. Half media changes were done every 3-4 days. 4 weeks after plating, cortical neurons were treated with DMSO, LCTB-21, or iso-LCTB-21, in NDM for 24 hrs. For western blot and kinase activity analysis: media was removed and 1x dPBS was added to each well. Cells were scraped off of the well and transferred to an Eppendorf tube. Cells were pelleted and stored at -80°C until analysis. For immunocytochemistry: media was removed and cells were fixed in 4% PFA for 15 minutes. Neurons were washed and stored in 1x PBS until analysis.

### Cell Viability CellTiter-Glo® 2.0 Assay

NPCs and neurons were plated at a density of 50,000 cells per well in a 96 well plate. 24 hrs after plating, cells were treated with DMSO, LCTB-21, or iso-LCTB-21. 24 hrs after treatment, media was replaced, and the CellTiter-Glo® 2.0 Assay (Promega) was performed according to the manufacturer’s protocol. Luminescence was recorded with the Promega GlomMax Multi Detection System.

### Western Blots

Cell pellets were lysed on ice in homogenization buffer (25 mM MOPS (Sigma), 15 mM EGTA (Sigma), 15 mM MgCl2 (Sigma), 60 mM β-glycerophosphate (Sigma), 15 mM p-nitrophenylphosphate (pNPP, Sigma), 2 mM DL-dithiothreitol (DTT, Sigma), 1 mM Na_3_VO_4_ (Sigma), 1 mM NaF (Sigma), 1 mM di-sodium phenylphosphate (Sigma), protease inhibitor cocktail (Complete, Roche), pH 7,2) supplemented with 0.1% Nonidet P-40, and then centrifuged (17,000 g for 1 min at 4°C). Protein extracts were mixed (1:1 v/v) with sample buffer (2x NuPAGE LDS sample buffer, 200 mM DTT). Following heat denaturation, 25 µg of proteins were loaded on NuPAGE precast 4-12% Bis-Tris protein gels (ThermoFisher). Electrophoresis was performed in MOPS buffer. Rapid blot transfers were carried out at 2.5 A/25 V for 7 min. Membranes were blocked in milk (5% Regilait in Tris Buffered Saline with 0.1% Tween (TBST)) for 1 h. Membranes were then incubated with primary antibodies (Supporting Table 2). Finally, membranes were incubated for 1 h at RT with HRP-conjugated secondary antibodies (Supporting Table 2), and chemiluminescent detection was achieved with homemade ECL-Tris buffer (100 mM Tris pH 8.5, 0.009% H_2_O_2_, 0.225 mM p-coumaric acid, 1.25 mM luminol) with Fusion Fx7 camera software. Bands were quantified using ImageJ. Experiments were repeated at least three times.

### Kinase Activity Assay

Dosage of DYRK1A activity was performed on cell lysates using HPLC as previously described (Bui et al., 2014). To assess the activity of DYRK1A by HPLC, a Dansyl-conjugated peptide was designed, Dansyl–KKISGRLSPIMTEQ-NH_2_ (Dan–peptide), the sequence of which is derived from the human forkhead transcription factor FKHR. This transcription factor is known to be phosphorylated by DYRK1A on the Ser329 residue of the GRLSPIM motif. This peptide and its Ser329 phosphorylated product should be easily separated by reverse-phase HPLC and specifically quantified by detection of the fluorescence of the Dansyl moiety. The purity and identity of the fluorescein-labeled peptide substrate (Dan–peptide) was initially assessed by reverse-phase HPLC (Prominence Shimadzu UFLC [ultra-fast liquid chromatography] system interfaced with LabSolutions software). Samples were injected into a Nucleodur C18 column (length = 150 mm, internal diameter = 4.6 mm, particle size = 5 µm) at 45 °C. The mobile phase used for the separation consisted of two eluents; solvent A was water with 0.1% trifluoroacetic acid (TFA), and solvent B was acetonitrile with 0.1% TFA. Compounds were separated by an isocratic flow (85% A/15% B) rate of 1.0 ml/min. The products were monitored by fluorescence emission (l = 537 nm) after excitation at l = 375 nm and quantified by integration of the peak absorbance area, employing a calibration curve established with various known concentrations of peptides. Incubation of DYRK1A with the Dan–peptide in the presence of ATP leads to two peaks corresponding to the FAM–peptide (retention time of 5.5 min) and its phosphorylated form (retention time of 4.8 min). As expected, the amount of phosphorylated Dan–peptide calculated by integration of the peak (area under the curve, AUC) increases linearly with the time of incubation. Assays were performed in a 96-well ELISA plate in a total volume of 50 µL consisting of kinase buffer (50 mM Tris–HCl, 10 mM DTT, and 5 mM MgCl_2_), ATP (up to 1,000 µM), Dan–peptide substrate (up to 100 µM), and purified DYRK1A–DC (up to 0.5 ng) or cell extracts (25 µg total protein extract). Briefly, samples containing the enzyme were preincubated with peptide substrate at 37 °C for 1 min, and the reaction was started by the addition of ATP. At different time points (up to 30 min), 50 µL of HClO_4_ (15% in water) was added to stop the reaction, and 20 µL was automatically injected into the HPLC column.

### Immunocytochemistry

After treatment, cells were fixed in 4% PFA for 15 minutes. Cells were blocked with 5% Normal Donkey Serum (NDS) and permeabilized with 0.2% TX-100 in 1x PBS for 15 minutes at room temperature. Primary antibodies at desired concentrations (Supporting Table 2) with 5% NDS in 1x PBS were applied overnight at 4°C. Primary antibody solutions were washed off with 1x PBS. Secondary antibodies at 1:500 with 5% NDS in 1x PBS were applied for 30 minutes at room temperature. Cells were washed with 1x PBS and stained with 1:100 Hoechst 33342 for 5 minutes at room temperature. Cells were washed with 1x PBS and mounted with Fluoromount-G™ (Invitrogen).

### High Content Imaging Analysis

NPCs and neurons were plated on 96-well plates with #1.5 bottom (Cellvis). Proliferation and cell culture composition analyses were carried out with the Nano High-Content Imaging System (Molecular Devices) with the MetaXpress software. 4-6 wells per treatment condition were acquired. NPC images were acquired at 20x with 9 sites/well. Neuron images were acquired at 20x with 16 sites/well. Representative images were acquired at 20x or 40x using the Nikon Ti2 Eclipse microscope.

## Supporting Tables

**Supporting Table 1.**
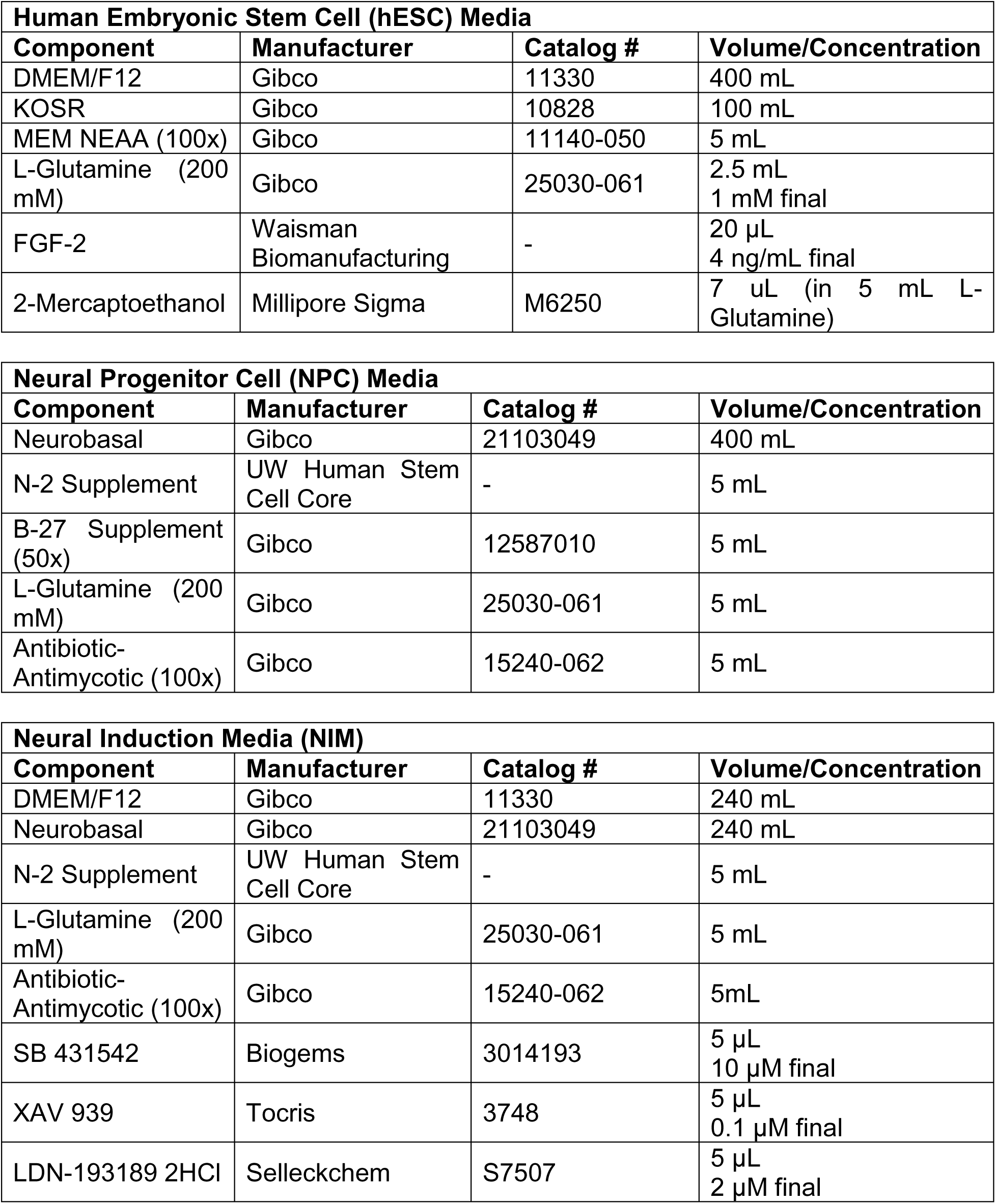
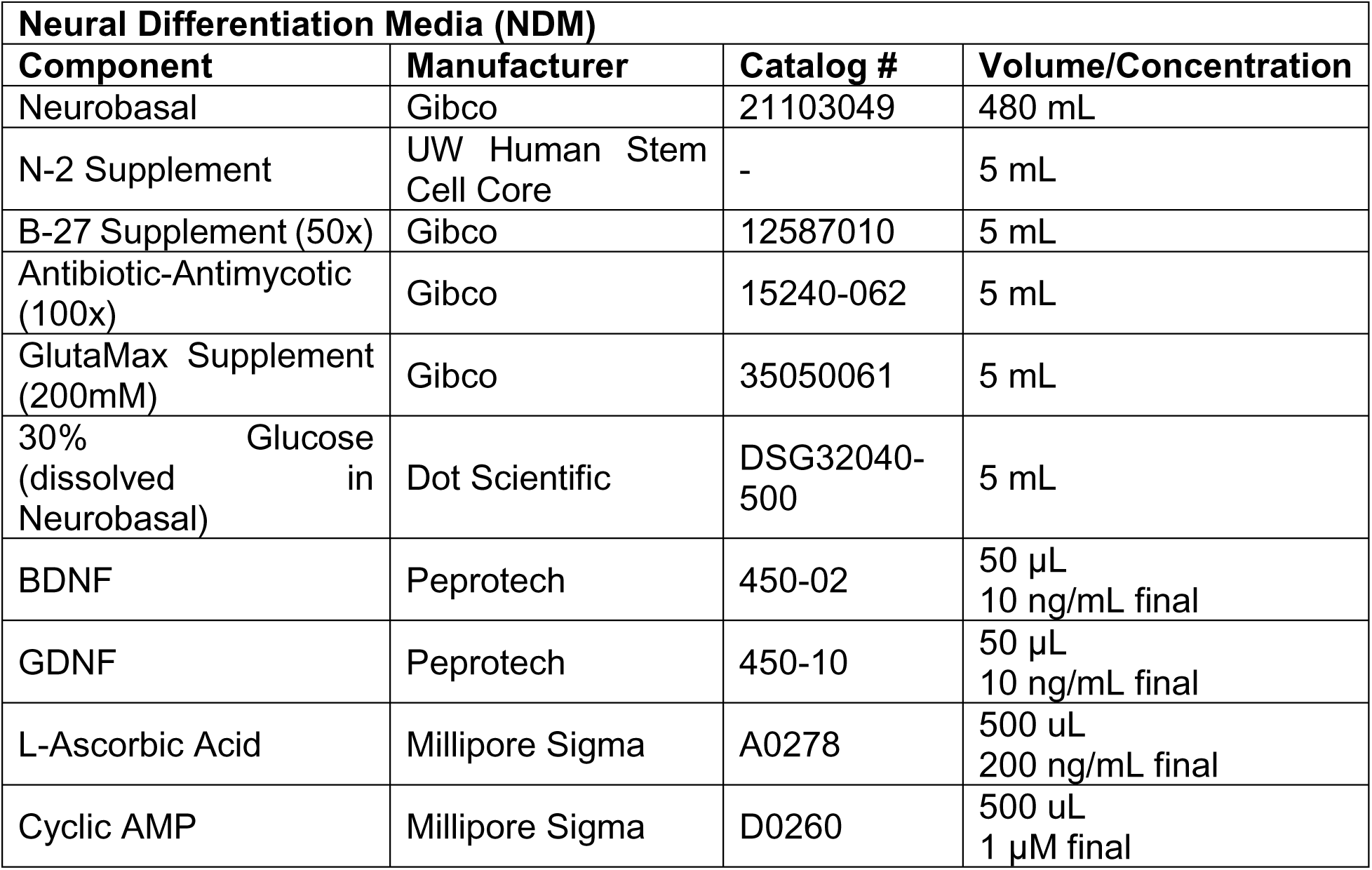
Media Recipes and Components.

**Supporting Table 2.**
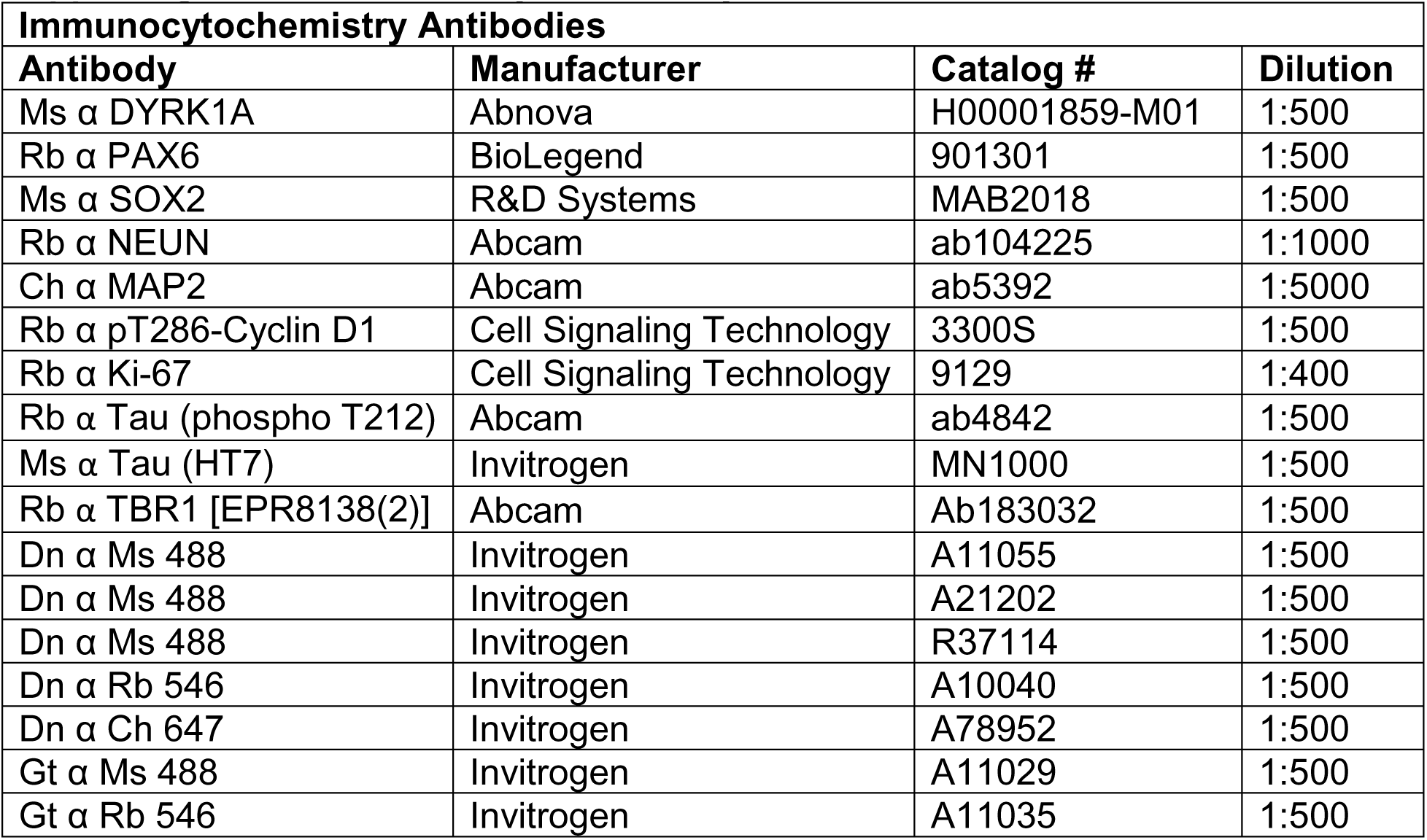
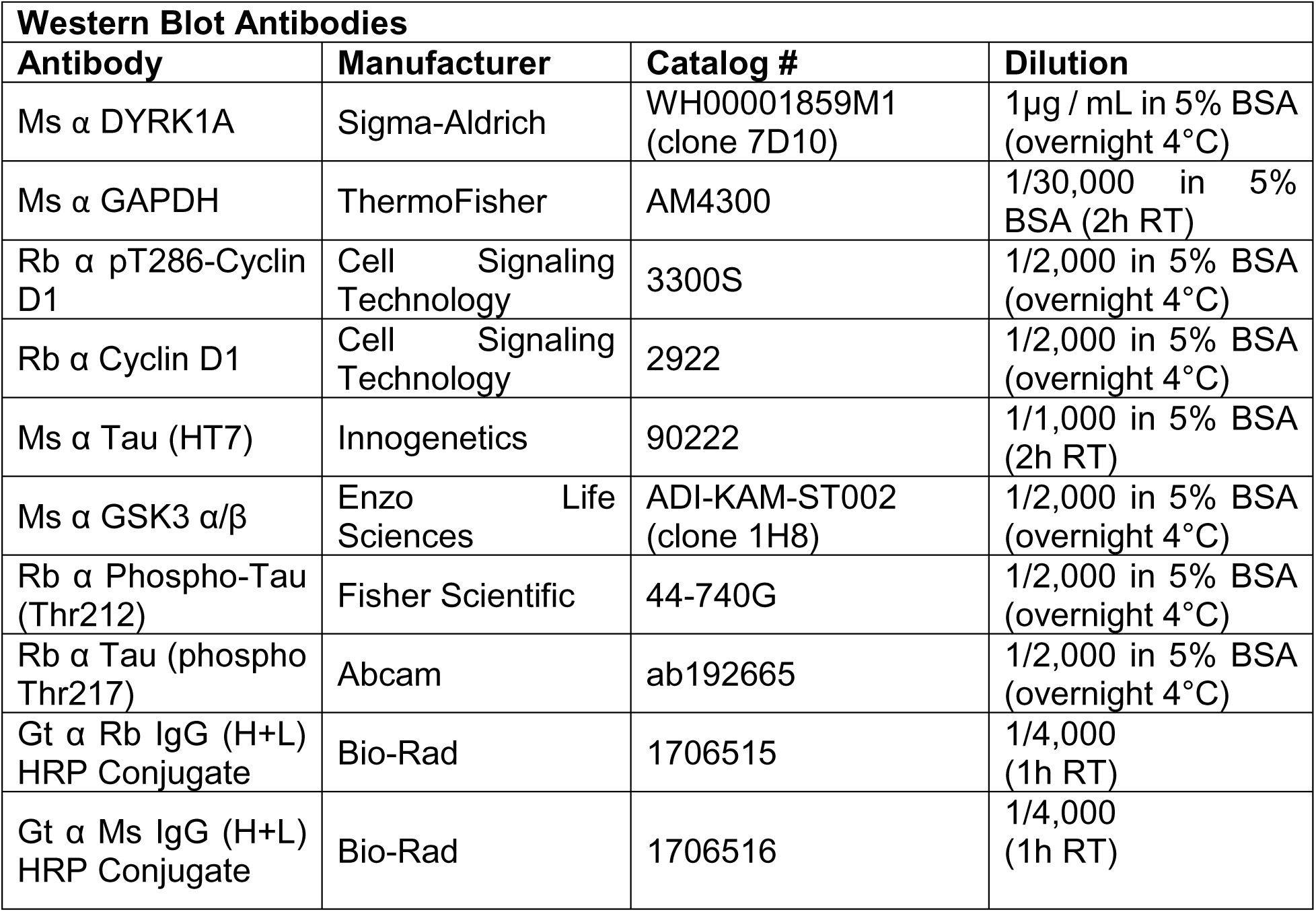
Immunocytochemistry and Western Blot Antibodies.

| **Immunocytochemistry Antibodies** |  |  |  |
| --- | --- | --- | --- |
| **Antibody** | **Manufacturer** | **Catalog #** | **Dilution** |
| Ms $\alpha$ DYRK1A | Abnova | H00001859-M01 | 1:500 |
| Rb $\alpha$ PAX6 | BioLegend | 901301 | 1:500 |
| Ms $\alpha$ SOX2 | R&D Systems | MAB2018 | 1:500 |
| Rb $\alpha$ NEUN | Abcam | ab104225 | 1:1000 |
| Ch $\alpha$ MAP2 | Abcam | ab5392 | 1:5000 |
| Rb $\alpha$ pT286-Cyclin D1 | Cell Signaling Technology | 3300S | 1:500 |
| Rb $\alpha$ Ki-67 | Cell Signaling Technology | 9129 | 1:400 |
| Rb $\alpha$ Tau (phospho T212) | Abcam | ab4842 | 1:500 |
| Ms $\alpha$ Tau (HT7) | Invitrogen | MN1000 | 1:500 |
| Rb $\alpha$ TBR1 [EPR8138(2)] | Abcam | Ab183032 | 1:500 |
| Dn $\alpha$ Ms 488 | Invitrogen | A11055 | 1:500 |
| Dn $\alpha$ Ms 488 | Invitrogen | A21202 | 1:500 |
| Dn $\alpha$ Ms 488 | Invitrogen | R37114 | 1:500 |
| Dn $\alpha$ Rb 546 | Invitrogen | A10040 | 1:500 |
| Dn $\alpha$ Ch 647 | Invitrogen | A78952 | 1:500 |
| Gt $\alpha$ Ms 488 | Invitrogen | A11029 | 1:500 |
| Gt $\alpha$ Rb 546 | Invitrogen | A11035 | 1:500 |

| **Western Blot Antibodies** |  |  |  |
| --- | --- | --- | --- |
| **Antibody** | **Manufacturer** | **Catalog #** | **Dilution** |
| Ms $\alpha$ DYRK1A | Sigma-Aldrich | WH00001859M1<br>(clone 7D10) | 1μg / mL in 5% BSA<br>(overnight 4°C) |
| Ms $\alpha$ GAPDH | ThermoFisher | AM4300 | 1/30,000 in 5%<br>BSA (2h RT) |
| Rb $\alpha$ pT286-Cyclin<br>D1 | Cell Signaling<br>Technology | 3300S | 1/2,000 in 5% BSA<br>(overnight 4°C) |
| Rb $\alpha$ Cyclin D1 | Cell Signaling<br>Technology | 2922 | 1/2,000 in 5% BSA<br>(overnight 4°C) |
| Ms $\alpha$ Tau (HT7) | Innogenetics | 90222 | 1/1,000 in 5% BSA<br>(2h RT) |
| Ms $\alpha$ GSK3 $\alpha/\beta$ | Enzo Life<br>Sciences | ADI-KAM-ST002<br>(clone 1H8) | 1/2,000 in 5% BSA<br>(overnight 4°C) |
| Rb $\alpha$ Phospho-Tau<br>(Thr212) | Fisher Scientific | 44-740G | 1/2,000 in 5% BSA<br>(overnight 4°C) |
| Rb $\alpha$ Tau (phospho<br>Thr217) | Abcam | ab192665 | 1/2,000 in 5% BSA<br>(overnight 4°C) |
| Gt $\alpha$ Rb IgG (H+L)<br>HRP Conjugate | Bio-Rad | 1706515 | 1/4,000<br>(1h RT) |
| Gt $\alpha$ Ms IgG (H+L)<br>HRP Conjugate | Bio-Rad | 1706516 | 1/4,000<br>(1h RT) |

## Abbreviations and nomenclature

AD: Alzheimer’s disease
DS: Down syndrome
DYRK1A: Dual-specificity tyrosine phosphorylation-regulated kinase 1A
Hsa21: Human chromosome 21
iPSC: Induced pluripotent stem cell
iso-LCTB-21: iso -Leucettinib-21 (inactive isomer)
LCTB-21: Leucettinib-21
NPC: Neural progenitor cell
T21: Trisomy 21

---

**Supporting Figure 1.**
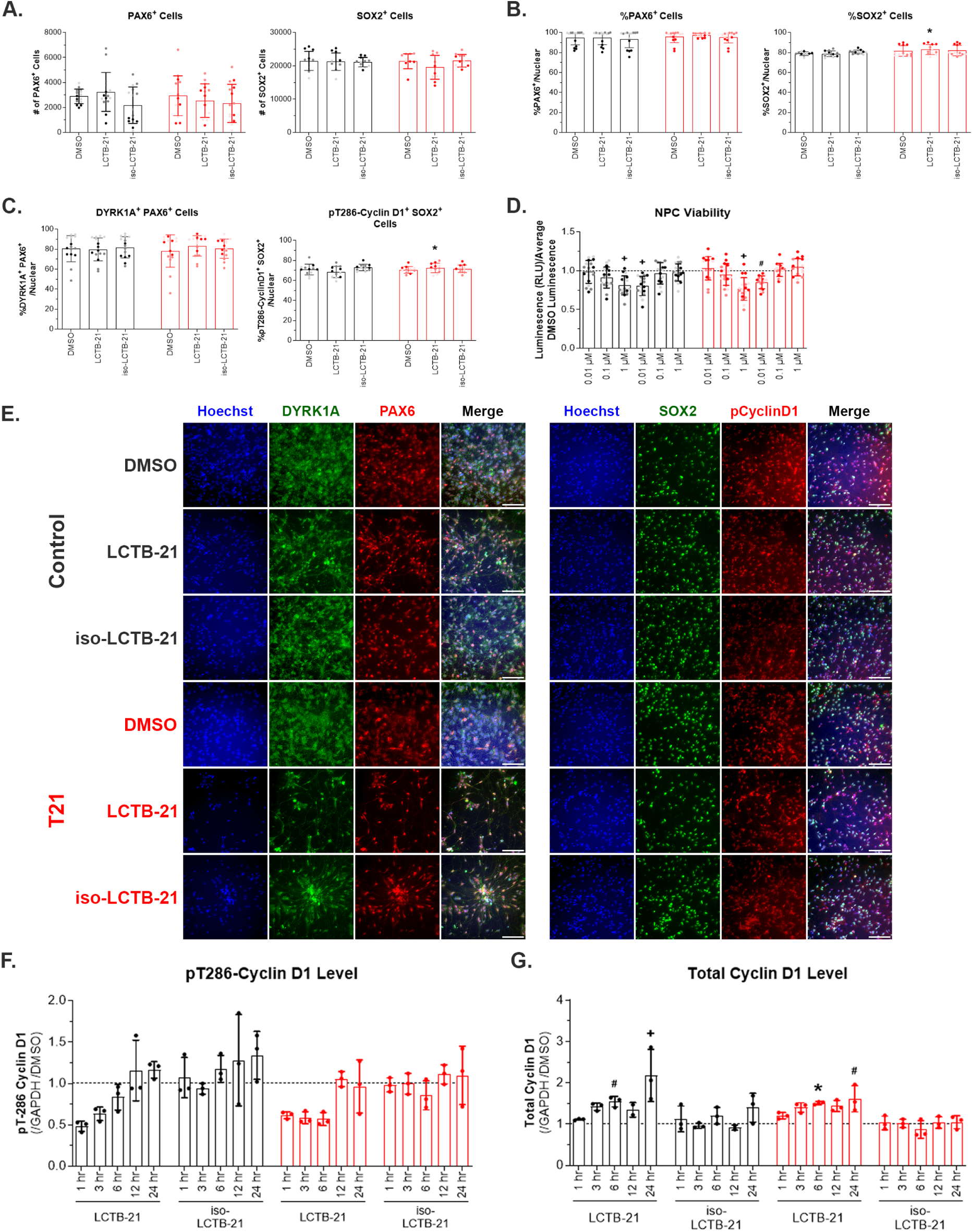
A) Pax6+ and Sox2+ progenitor cells in culture. Treatment with LCTB-21 has no effect on the number of NPCs in culture. B) Percent of Pax6+ and Sox2+ cells in culture. Treatment with LCTB-21 has no effect on the percentage of NPCs in culture relative to respective DMSO. There is a slight increase in LCTB-21 treated T21 NPCs compared to LCTB-21 treated control NPCs. C) Percent of DYRK1A expressing Pax6+ cells and pT286-cyclin D1 expressing Sox2+ cells. Treatment with LCTB-21 has no effect on the percentage of DYRK1A expressing or pT286-cyclin D1 expressing NPCs in culture. There is a slight increase in pT286-cyclin D1+Sox2+ T21 NPCs treated with LCTB-21 compared to LCTB-21 treated control NPCs. D) NPC viability. There is a slight decrease in the viability of NPCs treated with 1 μM LCTB-21 and 0.01 μM iso-LCTB-21 relative to respective DMSO (represented by the dashed line). E) Representative images of DYRK1A+ Pax6+ NPCs and pT286-cyclin D1+ Sox2+ NPCs (20x objective, 100 μm scale bar). F) pT286-cyclin D1 levels over time. pT286-cyclin D1 is reduced in control and T21 NPCs within 1 hour of treatment with LCTB-21 and this reduction persists for 6 hours compared to respective DMSO treatment (represented by the dashed line). G) Total cyclin D1 levels over time. Total cyclin D1 increases in control and T21 NPCs within 6 hours after treatment with LCTB-21 compared to respective DMSO treatment (represented by the dashed line). Data presented as mean + SD. P-values from Two-way ANOVA with post-hoc tests are presented as * ≤ 0.05, # ≤ 0.01, X ≤ 0.001, + ≤ 0.0001. Two-way ANOVA results, post-hoc tests, and P-values, and excluded values are listed in Supporting Table 3.

**Supporting Figure 2.**
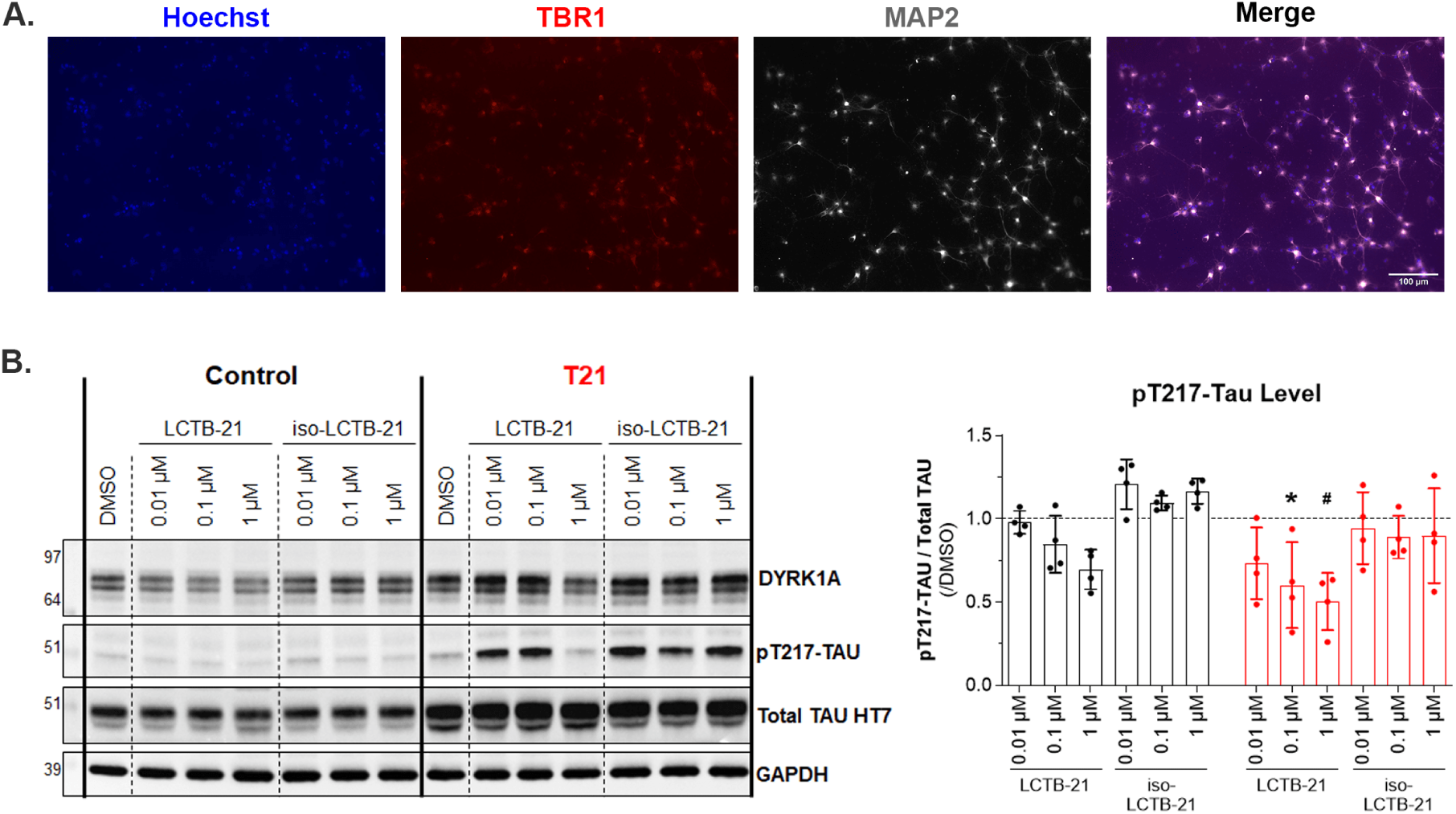
A) TBR1 expression in neurons, confirming the differentiation of iPSCs into cortical neurons (20x objective, 100 μm scale bar). B) Western blot and quantification of Total Tau and pT217 Tau levels in control and T21 cortical neurons. pT217 Tau levels decrease as DYRK1A activity decreases with increasing doses of LCTB-21. Data presented as mean + SD. P-values from Two-way ANOVA with post-hoc tests are presented as * ≤ 0.05, # ≤ 0.01, X ≤ 0.001, + ≤ 0.0001. Two-way ANOVA results, post-hoc tests, and P-values, and excluded values are listed in Supporting Table 3.

